## Supplemental Information including extended methods and additional results for "Resource landscape shapes the composition and stability of the human vaginal microbiota" (from another version of this preprint: version 5, 2025-09-07)


#### 1 Extended methods

##### 1.1 Main text model

Our vaginal microbiota model is described by the following set of equations:

$$\frac{\mathrm{d}C_1}{\mathrm{d}t} = -\gamma_C\ C_1 + \frac{\lambda_1\ R_s\ C_1}{\kappa + R_s} - \alpha_{1,1}\ C_1\ C_1 \tag{1a}$$

$$\frac{\mathrm{d}C_4}{\mathrm{d}t} = -\gamma_C\ C_4 + \frac{\lambda_4\ (R_p + R_s)\ C_4}{\kappa + R_p + R_s} - \alpha_{1,4}\ C_1\ C_4 \tag{1b}$$

$$\frac{\mathrm{d}R_p}{\mathrm{d}t} = \eta_{Rp}\frac{C_4}{\theta + C_4} - \gamma_R\ R_p - \frac{\lambda_4\ R_p\ C_4}{\epsilon(\kappa + R_p + R_s)} \tag{1c}$$

$$\frac{\mathrm{d}R_s}{\mathrm{d}t} = \eta_{R_s} - \gamma_R\ R_s - \frac{\lambda_1\ R_s\ C_1}{\epsilon(\kappa + R_s)} - \frac{\lambda_4\ R_s\ C_4}{\epsilon(\kappa + R_p + R_s)} \tag{1d}$$

The subscripts 1 and 4 refer to CST I and CST IV bacteria and the subscripts $p$ and $s$ refer to private and shared resources, respectively.

##### 1.2 Model restructuring

To facilitate model exploration, we first restructure the above model (eq. 1) into equations tracking the dynamics of the total bacterial ($C = C_1 + C_4$) and resource ($R = R_p + R_s$) densities and proportions that make up the CST IV bacteria ($q = \frac{C_4}{C}$) and their private resource ($p = \frac{R_p}{R}$). We then introduce $\chi$, the proportion of newly supplied resource that is private to $C_4$ (such that $\eta_{R_p} = \chi\ \eta_R$ and $\eta_{R_s} = (1 - \chi)\ \eta_R$ where $\eta_R$ is the overall supply). We also define $\psi$ as the ratio of CST IV inhibition and self-inhibition (i.e., $\frac{\alpha_{1,4}}{\alpha_{1,1}}$).

We then set the time scale of the system relative to the CST I growth rate ($\lambda_1$) such that selected parameters become relative to it and variables are dimensionless (i.e., non-dimensionalisation). Next, we assume a separation of timescales between consumers and resources and solve (owing to the Tikhonov-Fenichel theorem) for quasi-equilibrium conditions for resources such that the resource availability is held constant for a given density of consumers. In doing so, we now only explicitly track the consumer dynamics.

**Rearranging as total densities and proportions:** The dynamics of the total bacterial density is the sum of Eq. 1a and 1b:

$$\frac{\mathrm{d}C}{\mathrm{d}t} = \frac{\mathrm{d}C_1}{\mathrm{d}t} + \frac{\mathrm{d}C_4}{\mathrm{d}t} \tag{2a}$$

$$= \left(-\gamma_C - \alpha_{1,1}C(1-q)(1-(1-\psi)q) - \frac{\lambda_1 R(-p\ q + p + q - 1)}{\kappa - p\ R + R} + \frac{\lambda_4\ q\ R}{\kappa + R}\right) C. \tag{2b}$$

Table 1: Mechanistic model parameter notations, descriptions and values (range explored are shown inside square brackets). Rates are in units per day. Control parameters explored in sensitivity analyses (Supplementary Text S2) are marked with an asterisk. No value is listed for $\lambda_1$ as it is factored out.

| Symbol | Description | Value [Range] |
|---|---|---|
| Bacteria | | |
| $\gamma_C$ | Background clearance rate of bacteria | $\lambda_1 \ \overline{\gamma_C}$ |
| $\overline{\gamma_C}$ | Background clearance rate of bacteria relative to $\lambda_1$ | 0.1 [0.01, 0.454]* |
| $\kappa$ | Half-saturation constant for bacterial growth | 1 |
| $\theta$ | Half-saturation constant for sialic acid cleaving | 0.112 [0.01,1]* |
| $\lambda_1$ | Intrinsic growth rate of CST I bacteria | |
| $\lambda_4$ | Intrinsic growth rate of CST IV bacteria | $\lambda_1 \ \overline{\lambda_4}$ |
| $\overline{\lambda_4}$ | Ratio of $\lambda_4$ to $\lambda_1$ | 0.870 [0.622, 1.440]* |
| $\alpha_{1,1}$ | Self-inhibition rate of CST I | $\lambda_1 \ \overline{\alpha_{1,1}}$ |
| $\overline{\alpha_{1,1}}$ | Self-inhibition rate of CST I relative to $\lambda_1$ | 0.483 [0.01, 0.618]* |
| $\alpha_{1,4}$ | Inhibition rate of CST IV by CST I | $\psi \ \alpha_{1,1}$ |
| $\psi$ | Ratio of $\alpha_{1,4}$ to $\alpha_{1,1}$ | 2 [1.01, 5]* |
| Resources | | |
| $\eta_{R_p}$ | Supply rate for CST IV private resource | $\eta_R \ \chi$ |
| $\eta_{R_s}$ | Supply rate for shared resource | $\eta_R(1-\chi)$ |
| $\eta_R$ | Overall resource supply rate | $\lambda_1 \ \overline{\eta_R}$ |
| $\overline{\eta_R}$ | Overall resource supply rate relative to $\lambda_1$ | $\overline{\tau}/\epsilon$ |
| $\chi$ | Proportion of resource supply that is private to CST IV | $4.414e^{-3}$ [0,1]* |
| $\gamma_R$ | Background clearance rate of resources | $\lambda_1 \ \overline{\gamma_R}$ |
| $\overline{\gamma_R}$ | Background clearance rate of resources relative to $\lambda_1$ | 0 |
| $\epsilon$ | Yield per unit of resource | $\overline{\tau}/\overline{\eta_R}$ |
| $\overline{\tau}$ | Productivity: product of yield ($\epsilon$) and time-scaled supply ($\overline{\eta_R}$) | 0.466 [0.1, 0.9]* |

Next, we derive $\frac{dq}{dt}$. Substituting $\frac{dC}{dt}$ and $\frac{dC_4}{dt}$ (Eq. 2 and 1b), we get:

$$\frac{dq}{dt} = \frac{d}{dt}\frac{C_4}{C} = \frac{\frac{dC_4}{dt}C - \frac{dC}{dt}C_4}{C^2} = \frac{\frac{dC_4}{dt} - \frac{dC}{dt}q}{C} \tag{3a}$$

$$= \left(\alpha_{1,1}C(1-q)(1-\psi) - \frac{\lambda_1(1-p)R}{\kappa - pR + R} + \frac{\lambda_4 \ R}{\kappa + R}\right)(1-q)q. \tag{3b}$$

Similarly, the dynamics of the overall resource is simply the sum of Eq. 1c and 1d.

$$\frac{dR}{dt} = \frac{dR_p}{dt} + \frac{dR_s}{dt} \tag{4a}$$

$$= \frac{(\kappa+R)\left(C\lambda_1(1-p)(1-q) + \gamma_R\epsilon(\kappa - pR + R)\right) + C\lambda_4 q(\kappa - pR + R)}{\epsilon(\kappa+R)(-(1-p)R-\kappa)}R - \frac{\eta_R\theta\chi}{C \ q + \theta} + \eta_R. \tag{4b}$$

Next, we define $\frac{dp}{dt}$ and substituting $\frac{dR}{dt}$ and $\frac{dR_p}{dt}$ (Eq. 4b and 1c):

$$\frac{dp}{dt} = \frac{d}{dt}\frac{R_p}{R} = \frac{\frac{dR_p}{dt}R - \frac{dR}{dt}R_p}{R^2} = \frac{\frac{dR_p}{dt} - \frac{dR}{dt}p}{R} \tag{5a}$$

$$= \frac{\eta_R \ \chi(C \ q + \theta \ p)}{R(C \ q + \theta)} + \frac{\lambda_1(1-p)p(1-q)C}{\epsilon(\kappa - p \ R + R)} - \frac{\eta_R \ p}{R}. \tag{5b}$$

**Non-dimensionalisation:**
We rescale time according to $\lambda_1$. Rearranging the systems with respect to the dimensionless variable, $\bar{t}$

and setting $t_c = \frac{1}{\lambda_1}$ allows us to express the re-scaled system as:

$$\frac{\mathrm{d}C}{\mathrm{d}\bar{t}} = \left(\overline{\alpha_{1,1}}C(1-q)(q(1-\psi)-1) - \overline{\gamma_C} + \frac{(1-p)(1-q)R}{\kappa - p\,R + R} + \frac{\overline{\lambda_4}qR}{\kappa + R}\right)C \tag{6a}$$

$$\frac{\mathrm{d}q}{\mathrm{d}\bar{t}} = \left(\overline{\alpha_{1,1}}C(1-q)(1-\psi) - \frac{(1-p)R}{\kappa - p\,R + R} + \frac{\overline{\lambda_4}R}{\kappa + R}\right)(1-q)q \tag{6b}$$

$$\frac{\mathrm{d}R}{\mathrm{d}\bar{t}} = \overline{\eta_R} - \frac{\overline{\eta_R}\theta\chi}{Cq+\theta} + \frac{R(\kappa+R)(C(1-p)(1-q) + \overline{\gamma_R}\epsilon(\kappa - p\,R + R)) + C\overline{\lambda_4}qR(\kappa - pR + R)}{\epsilon(\kappa+R)(-(1-p)R - \kappa)} \tag{6c}$$

$$\frac{\mathrm{d}p}{\mathrm{d}\bar{t}} = \frac{\overline{\eta_R}\chi(Cq + \theta\,p)}{R(Cq+\theta)} + \frac{C(1-p)p(1-q)}{\epsilon(\kappa - p\,R + R)} - \frac{\overline{\eta_R}\,p}{R}. \tag{6d}$$

where $^-$ on parameters indicate scaling by $\lambda_1$ such that $\overline{\eta_R} = \frac{\eta_R}{\lambda_1}$, $\overline{\gamma_R} = \frac{\gamma_R}{\lambda_1}$, $\overline{\lambda_4} = \frac{\lambda_4}{\lambda_1}$, $\overline{\alpha_{1,1}} = \frac{\alpha_{1,1}}{\lambda_1}$ and $\overline{\gamma_C} = \frac{\gamma_C}{\lambda_1}$.

**Resource quasi-equilibrium and final consumer system:** While resource concentrations can change rapidly due to microbial uptake, the assimilation of those resources into biomass (i.e., bacterial growth) typically occurs on a slower timescale due to intracellular metabolic processing and regulation. By setting $\frac{\mathrm{d}R}{\mathrm{d}\bar{t}} = 0$ and $\frac{\mathrm{d}p}{\mathrm{d}\bar{t}} = 0$, we reflect the fast dynamics of resource consumption relative to bacterial growth. Within this quasi-instantaneous timescale, we assume that spontaneous decay and outflow of resources are negligible compared to bacterial uptake (i.e., $\overline{\gamma_R} = 0$), such that resource loss is driven almost entirely by consumption. This separation of time scales allows us to solve for quasi-equilibrium values $\hat{R}$ and $\hat{p}$, capturing the rapid reshaping of resource pools by bacteria while allowing bacterial biomass to evolve on a slower timescale. This approach is consistent with growth models of microorganisms that distinguish between fast uptake and slower growth processes (e.g., (Droop, 1968)):

$$\hat{R} = \frac{\kappa\,\overline{\tau}\left(-\chi(C\,q+\theta)(\theta\,\overline{\lambda_4} + \overline{\tau} - (1-q)C) + \overline{\lambda_4}(C\,q+\theta)^2 + \theta\,\overline{\tau}\,\chi^2\right)}{-\overline{\tau}\,\chi(C\,q+\theta)(-C(1-\overline{\lambda_4})q + C - \theta\,\overline{\lambda_4} - \overline{\tau}) + \overline{\lambda_4}(C\,q+\theta)^2(-C(1-\overline{\lambda_4})q + C - \overline{\tau}) - \theta\,\overline{\tau}^2\chi^2} \tag{7a}$$

$$\hat{p} = \frac{C\,\chi(q\,\overline{\tau}\,\chi - (1-(1-\overline{\lambda_4})q)(C\,q+\theta))}{\chi\,(Cq+\theta)(\theta\,\overline{\lambda_4} + \overline{\tau} - (1-q)C) - \overline{\lambda_4}(C\,q+\theta)^2 - \theta\,\overline{\tau}\,\chi^2} \tag{7b}$$

where $\overline{\tau} = \epsilon\,\overline{\eta_R}$. Substituting resource variables ($R$ and $p$) in the consumer equations (Eq. 6a & b), with their quasi-equilibrium conditions ($\hat{R}$ and $\hat{p}$; Eq. 7), we obtain the final consumer system, which is governed by seven parameters: $\chi$, $\theta$, $\overline{\tau}$, $\overline{\lambda_4}$, $\overline{\gamma_C}$, $\overline{\alpha_{1,1}}$, $\psi$.

#### 1.3 Parameterising the mechanistic model

$\theta$, $\kappa$, $\chi$, $\overline{\tau}$: Solving for productivity ($\overline{\tau}$) and proportion of resource supply that is private to CST IV ($\chi$) in Eq. 7, we get:

$$\overline{\tau} = \frac{\overline{\lambda_4}\,\hat{R}(\hat{C}\,\hat{q} + \theta\,\hat{p})}{\kappa + \hat{R}} + \frac{\hat{C}(1-\hat{p})(1-\hat{q})\hat{R}}{\kappa - \hat{p}\,\hat{R} + \hat{R}} \tag{8a}$$

$$\chi = \frac{\overline{\lambda_4}\,\hat{p}(\hat{C}\,\hat{q} + \theta)(-(1-\hat{p})\hat{R} - \kappa)}{\overline{\lambda_4}(-(1-\hat{p})\hat{R} - \kappa)(\hat{C}\,\hat{q} + \theta\,\hat{p}) - \hat{C}(1-\hat{p})(1-\hat{q})(\kappa + \hat{R})} \tag{8b}$$

The right-hand side of these equations (Eq. 8) can be parameterised empirically, except for the half-saturation constants $\theta$ and $\kappa$ whose exact values are unknown. Thus, estimates of $\overline{\tau}$ and $\chi$ are conditioned upon $\theta$ and $\kappa$. To demonstrate the typical equilibrium behaviour of the model, we set default $\theta$ and $\kappa$ as the median CST IV bacteria density (0.1125) and median total resource metabolite quantity (i.e., 1) in the normalised scale, respectively. With these defaults, we obtain the median $\overline{\tau}$ and $\chi$ of 0.466 and $4.414 \times 10^{-3}$, respectively. We conduct sensitivity analyses to explore the effect of assuming a range of $\theta$ and $\kappa$ values (Supplementary Text S3).

$\overline{\lambda_4}$: The relative *in vivo* growth rates of *G. vaginalis* to *L. crispatus*, representing CST IV and I, respectively, were estimated leveraging coincident isolate genome sequences and vaginal shotgun metagenomes. For each species, five strains were isolated from women for which concurrent shotgun metagenome data were available (see Auxiliary Data Files). Sequence reads from the metagenomes were mapped back to the corresponding isolate's genome sequence using `Bowtie2` (Langmead and Salzberg, 2012) and then `iRep` was used to estimate the index of replication (Brown et al., 2016). These values can be thought of as the percentage of cells which are actively replicating their chromosome (e.g., a value of 1.5 would be 50% replicating). Since our interest here is to estimate the relative (as opposed to the absolute) growth rate between *G. vaginalis* and *L. crispatus*, we take advantage of the ratio of the index of replication, assuming comparable doubling times between the two species in vivo. Through bootstrapping with 10,000 iterations, we report the median relative growth rate of *G. vaginalis* to *L. crispatus* ($\overline{\lambda_4}$) of 0.870 and a 95% interval of 0.622 to 1.440 (Figure S1). Our estimates were consistent with previously reported values (Anukam and Reid, 2008; Lee et al., 2020). We set the default to the median and explored values of the 95% interval.

$\overline{\gamma_C}$, $\overline{\alpha_{1,1}}$: We parameterise the background clearance rate ($\overline{\gamma_C}$) and self-inhibition rate of CST I relative to its growth ($\overline{\alpha_{1,1}}$) considering a CST I dominant community (in the absence of CST IV). Dividing Eq 1a by $\lambda_1$ to scale $t$ and rate parameters relative to $\lambda_1$, we obtain a unitless equation:

$$\frac{\mathrm{d}C_1}{\mathrm{d}\bar{t}} = \left(\frac{R_s}{\kappa + R_s} - \overline{\gamma_C} - \overline{\alpha_{1,1}}\, C_1\right) C_1. \tag{9}$$

Again, assuming that the resource dynamics are faster than bacteria and hence $R_s$ is a constant, we obtain a CST IV-free quasi-equilibrium expression for CST I as:

$$\hat{C}_1 = \frac{\frac{R_s}{\kappa+R_s} - \overline{\gamma_C}}{\overline{\alpha_{1,1}}} \tag{10}$$

Rearranging this equation with respect to $\overline{\alpha_{1,1}}$, we get:

$$\overline{\alpha_{1,1}} = \frac{\frac{R_s}{\kappa+R_s} - \overline{\gamma_C}}{C_1} \tag{11}$$

Subsetting the clinical data for CST I-classified communities, we plugged in the median glycogen derivative quantity as $R_s$ (i.e., 0.832 in the normalised scale), the median DNA quantity of CST I bacteria as $C_1$ (i.e., 0.734 in the normalised scale), and median total resource availability as $\kappa$ (i.e., 1 in the normalised scale). Finally, while the exact value of $\overline{\gamma_C}$ is unknown, we deduce that the background clearance rate needs to be much smaller than the *Lactobacillus* growth rate (i.e., $\overline{\gamma_C} \ll 1$) to sustain none-zero bacterial population size: therefore, we set the default $\overline{\gamma_C} = 0.1$. We then obtain a default estimate of $\overline{\alpha_{1,1}} = 0.483$. We explored a range of $\overline{\gamma_C}$ and $\overline{\alpha_{1,1}}$ values with the maximum value of either parameter constrained to ensure that the other remains positive.

$\psi$: CST IV bacteria are more susceptible than CSI I to acidic conditions created by lactic acid and suppressed by anti-microbial compounds produced by CST I(Atassi et al., 2006; France et al., 2022). Thus, the ratio of CST IV inhibition by CST I to CST I self-regulation ($\psi$) is greater than 1. We set the default to 2 and explored a range of values between 1 and 5.

### 2 Local sensitivity of qualitative outcomes to model parameters

We explored how key features of the vaginal microbiota ecology modulate microbiota community transitions using local sensitivity analyses of model parameters (i.e., by varying a parameter at a time and holding others constant at the default value).

$\overline{\tau}$: The productivity of the vaginal bacterial system ($\overline{\tau}$) is a compound parameter that represents the product of the overall resource supply and yield. While productivity has a relatively small impact on the lower bound of bacterial community bistability, we found that increasing $\overline{\tau}$ reduces the region of CST IV dominance such that a higher proportion of CST IV private resources is required to trigger a transition to full CST IV dominance.

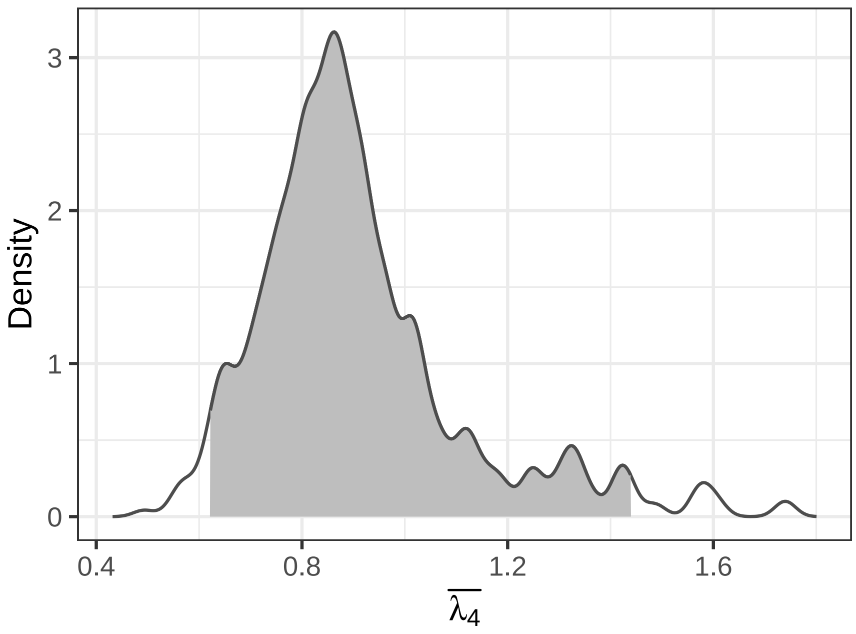


Figure S1: Distribution of bootstrapped $\overline{\lambda_4}$, i.e., relative growth rate of *Gardnerella vaginalis* to *Lactobacillus crispatus*. The grey shaded area indicates the 95% interval.

$\theta$: Sialic acid molecules are "havervested" or cleaved off by CST IV associated bacteria before consumption. The bacteria load required to achieve half-maximum sialic acid cleaving is expressed by $\theta$. We found that bacterial coexistence is only possible at low $\theta$ values and increasing $\theta$ tends to disfavour CST IV dominance.

$\overline{\lambda_4}$: We explored the role of the relative intrinsic growth rate of CST IV over CST I as the growth rate of both CST I-associated lactobacilli and CST IV-associated bacteria shows within-species variation across strains (Harwich et al., 2010; Van Der Veer et al., 2019) (Supplementary Text S1). We found that as $\overline{\lambda_4}$ approaches identity (i.e., CST I and CST IV have an equal growth rate), less CST IV private resource is needed to trigger a transition to CST IV dominance and maintain its dominance. Furthermore, the coexistence of the two communities becomes less likely as the relative CST IV growth rate increases.

$\overline{\gamma_C}$: The background bacterial clearance rate (i.e., Fig. S2: $\overline{\gamma_C}$) fluctuates during the menstrual cycle as vaginal discharge is particularly elevated during menses when blood and the functional layer of the uterine endometrium flow through the vagina canal. We find that coexistence is only possible at small $\overline{\gamma_C}$ values, while $\overline{\gamma_C}$ has a non-monotonic effect on the bistability window. Therefore, it is difficult to explain the association between menses and vaginal microbiota community transitions observed previously (Gajer et al., 2012) through our resource-driven perspective alone.

$\overline{\alpha_{1,1}}$: Lactic acid is considered lactobacilli's primary defence against BV-associated bacteria (O'Hanlon et al., 2013). In our model, $\overline{\alpha_{1,1}}$ represents the strength of CST I baseline regulation, which is known to be linked to lactic acid production and varies among *L. crispatus* strains (Argentini et al., 2022). We observed an increase in the propensity for CST I dominance with an increasing $\overline{\alpha_{1,1}}$ at the expense of coexistence between the two types. This finding is consistent with the established observation that acidic conditions are more favourable for the *Lactobacillus* growth (O'Hanlon et al., 2013). We also found that the upper boundary of bistability increases with $\overline{\alpha_{1,1}}$, such that a higher CST IV private resource ($\chi$) is required to trigger a transition to CST IV dominance in a highly acidic vaginal milieu. This finding offers a conceptual foundation for the use of lactic acid supplements as a preventative strategy against BV.

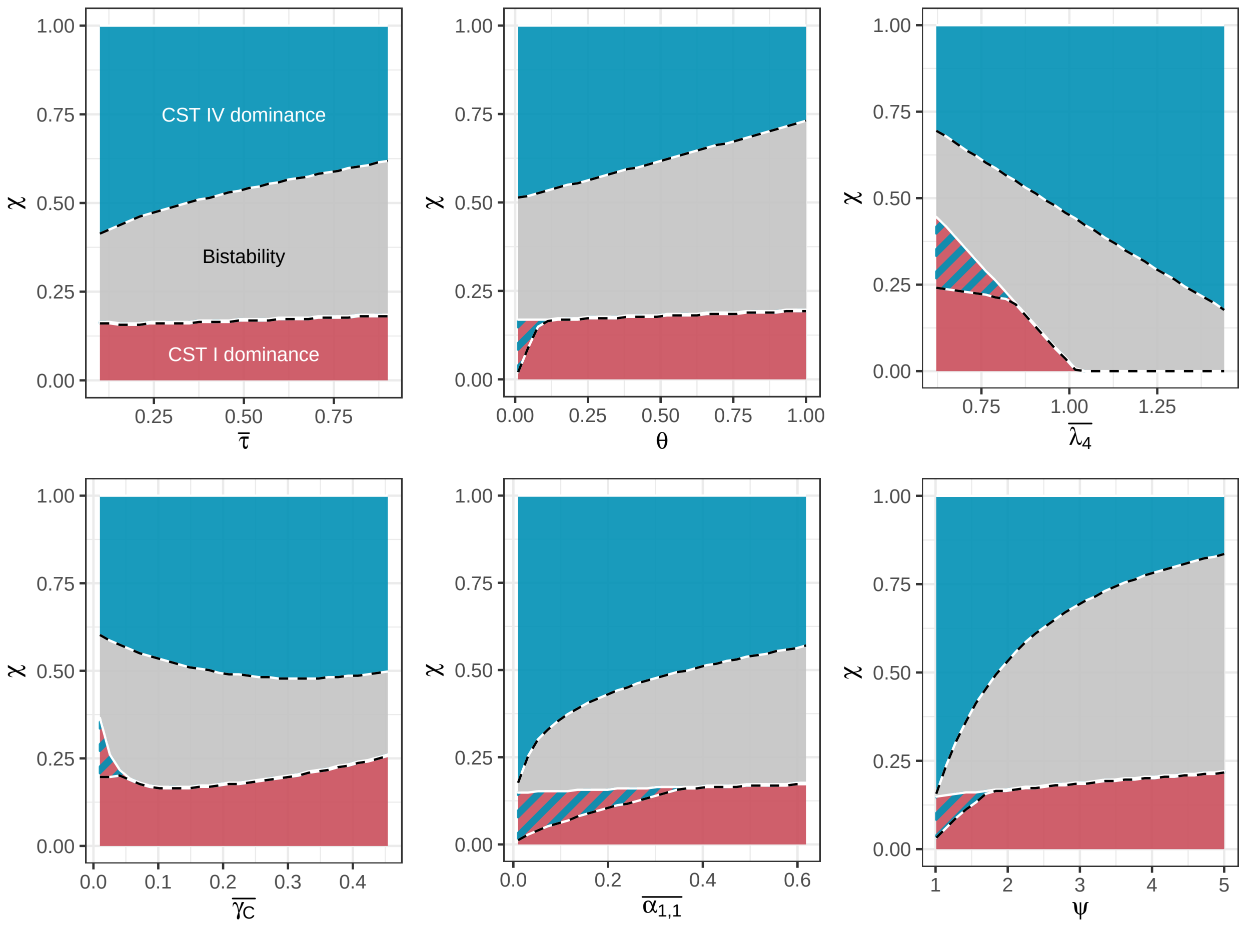


Figure S2: Vaginal bacterial community compositions as a function of control parameters ($\overline{\tau}$, $\theta$, $\overline{\lambda_4}$, $\overline{\gamma_C}$, $\overline{\alpha_{1,1}}$ and $\psi$) and the proportion of resource supply private to CST IV ($\chi$). The qualitative outcomes shown are monostable CST I dominance (red), CST IV dominance (blue), the coexistence of CST I and CST IV (red with blue stripes) and bistable hysteresis (light grey). The hysteresis window refers to the height of the grey band for a given parameter value on the x-axis. The areas of single community stability are demarcated by dashed lines. The minimum $q_0$ and $1 - q_0$ are set at 0.001, with an assumption that a small amount (0.1% of the total density) of less dominant bacteria-type seeds the vaginal milieu. The results are qualitatively identical across external input proportions as long as it maintains hysteresis (i.e., light grey shaded panels in Fig. 2 in the main text).

$\psi$: The strength of CST I-meditated population regulation against CST IV relative to self-regulation is denoted in our model by the parameter $\psi$ (Fig. S2). The qualitative impact of $\psi$ is similar to that of $\overline{\alpha_{1,1}}$: i.e., increasing $\psi$ increases the region of CST I dominance, requiring a higher proportion of CST IV-private resources to trigger a transition to CST IV.

### 3 Sensitivity to half-saturation constants, $\kappa$ and $\theta$

In our model the parameters $\kappa$ and $\theta$ signify the half-saturation constants for bacterial growth (i.e., resources required to achieve half the intrinsic growth rate) and for sialic acid cleaving (i.e., CST IV bacteria required to liberate half the sialic acid supplied as a bound source in the vaginal mucus). The default $\kappa$ and $\theta$ values were set to the median total resource metabolite quantity (i.e., 1) and median CST IV associated bacteria quantity (i.e., 0.1125) in the normalised scale — note that metabolite and bacteria quantities were normalised by the respective median. While these are plausible values, there is no specific empirical basis for them.

Estimating the half-saturation constants remains challenging as the values likely depend on microscale interactions that are unlikely to be accurately captured in well-mixed culture assays. However, realistic organ-on-a-chip systems — which better replicate spatial structure and physiological conditions — could offer a promising platform for systematically estimating the half-saturation constants across a range of biologically relevant contexts.

In our model exploration and data integration, the productivity ($\overline{\tau}$) and the proportion of resource supply private to CST IV ($\chi$) are key parameters conditional upon $\kappa$ and $\theta$ (Eq. 9). Thus, we graphically explored the sensitivity of $\overline{\tau}$ and $\chi$ to $\kappa$ and $\theta$ (Fig. S3). We found that the productivity ($\overline{\tau}$) estimate is largely influenced by the growth half-saturation constant ($\kappa$) such that the consumers become less productive as more resources are required for their growth (Fig. S3a). Conversely, we find that the estimate of the proportion of resource supply private to CST IV ($\chi$) is largely determined by the sialic acid cleaving half-saturation constant, $\theta$ (Fig. S3.b). If CST IV bacteria are inefficient at sialic acid cleaving (i.e., higher $\theta$), fewer unbound sialic acid molecules would be liberated per unit of bound sialic acid supplied to the system. Therefore, the lower the efficiency of sialic acid cleaving (i.e., higher $\theta$), the larger the initial supply of bound sialic acid (i.e., higher $\chi$) would have been to explain a given quantity of unbound sialic acid observed in the vaginal milieu. This sensitivity analysis indicates that absolute values of $\overline{\tau}$ and $\chi$ are not particularly informative unless $\kappa$ and $\theta$ can be more closely motivated from empirical data.

Nonetheless, our inference about the impact of resources through $\overline{\tau}$ and $\chi$ estimates would be qualitatively identical so long as the rank order of $\overline{\tau}$ and $\chi$ estimates are preserved regardless of the weakly informed $\kappa$ and $\theta$ values. To assess the extent to which $\kappa$ and $\theta$ influence the rank order of $\overline{\tau}$ and $\chi$ estimates, we estimated $\overline{\tau}$ and $\chi$ values for each data point (with microbiota and metabolomics data) assuming different $\kappa$ and $\theta$ values. We then calculated the Spearman's rank correlation coefficient between the estimates (i.e., $\overline{\tau}$ and $\chi$, separately) assuming the default and other plausible combinations of $\kappa$ and $\theta$ values. We observed that the rank orders of both $\overline{\tau}$ and $\chi$ are generally well-preserved with the correlation coefficient generally well over 0.7 (Fig. S3.b). For $\overline{\tau}$, relatively low correlations ($\sim 0.7$) were observed for low $\kappa$ (where growth requires fewer resources) and high $\theta$ (where sialic acid cleaving is inefficient) values (Fig. S3.b2). For $\chi$, we found observed relatively low correlations ($\sim 0.8$) at low (Fig. S3.b3) and high (Fig. S3.b4; where sialic acid cleaving is much more or less efficient than the assumed default, respectively).

Using clinical data and assuming default $\kappa$ and $\theta$ parameters, we demonstrated that relative and absolute supplies of resources private to CST IV communities are a major predictor of their dominance (Fig. 3 in the main text). To corroborate this finding across different combinations of half-saturation constants (i.e., $\kappa$ and $\theta$), we reproduced Fig. 3 from the main text with parameter combinations that produced the most divergent estimates of $\overline{\tau}$ and $\chi$ from the default combination (i.e., Fig. S3.b2, S3.b3, and S3.b4). We find that when we assume inefficient sialic acid cleaving (i.e., high $\theta$; Fig. S4.b2 & S4.b4), the impact of the proportion of resource supply private to CST IV ($\chi$) becomes weaker, whereas assuming efficient cleaving (i.e., low $\theta$; Fig. S4.b3) strengthens its influence. These findings are reasonable considering that the private resource supply would have little influence when the accessibility of private resources is limited and consumer dynamics are primarily influenced by shared resources. We find that the influence of absolute resource supplies is somewhat consistent regardless of the half-saturation constants assumed: CST I communities are always associated with low absolute private resource (i.e., sialic acid) supplies (Fig. S5).

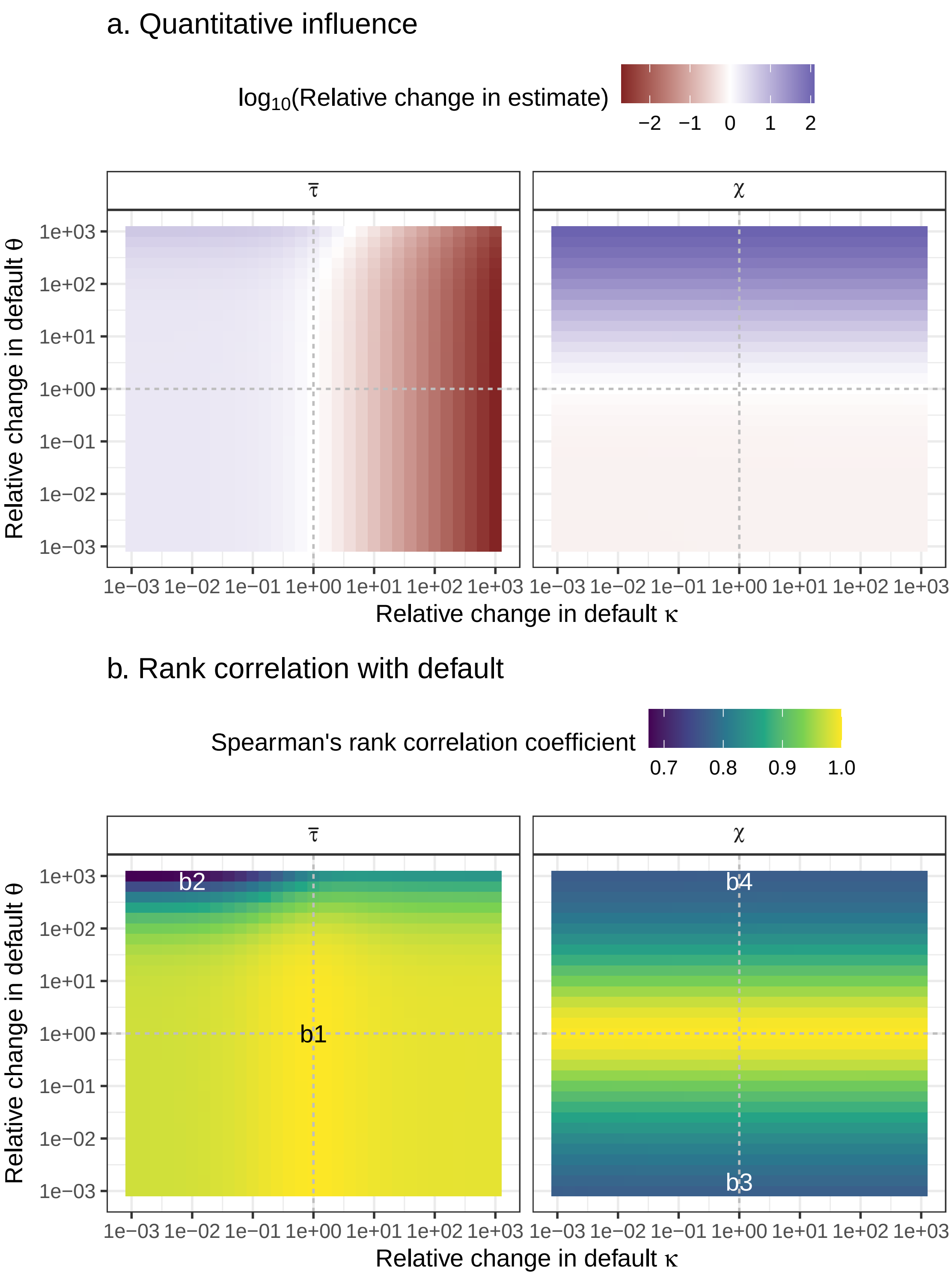


Figure S3: Sensitivity of the productivity ($\overline{\tau}$) and proportion of resource supply private to CST IV ($\chi$), to half-saturation constants $\kappa$ (x-axis) and $\theta$ (y-axis). b1 indicates the default parameter combination of $\kappa$ and $\theta$.

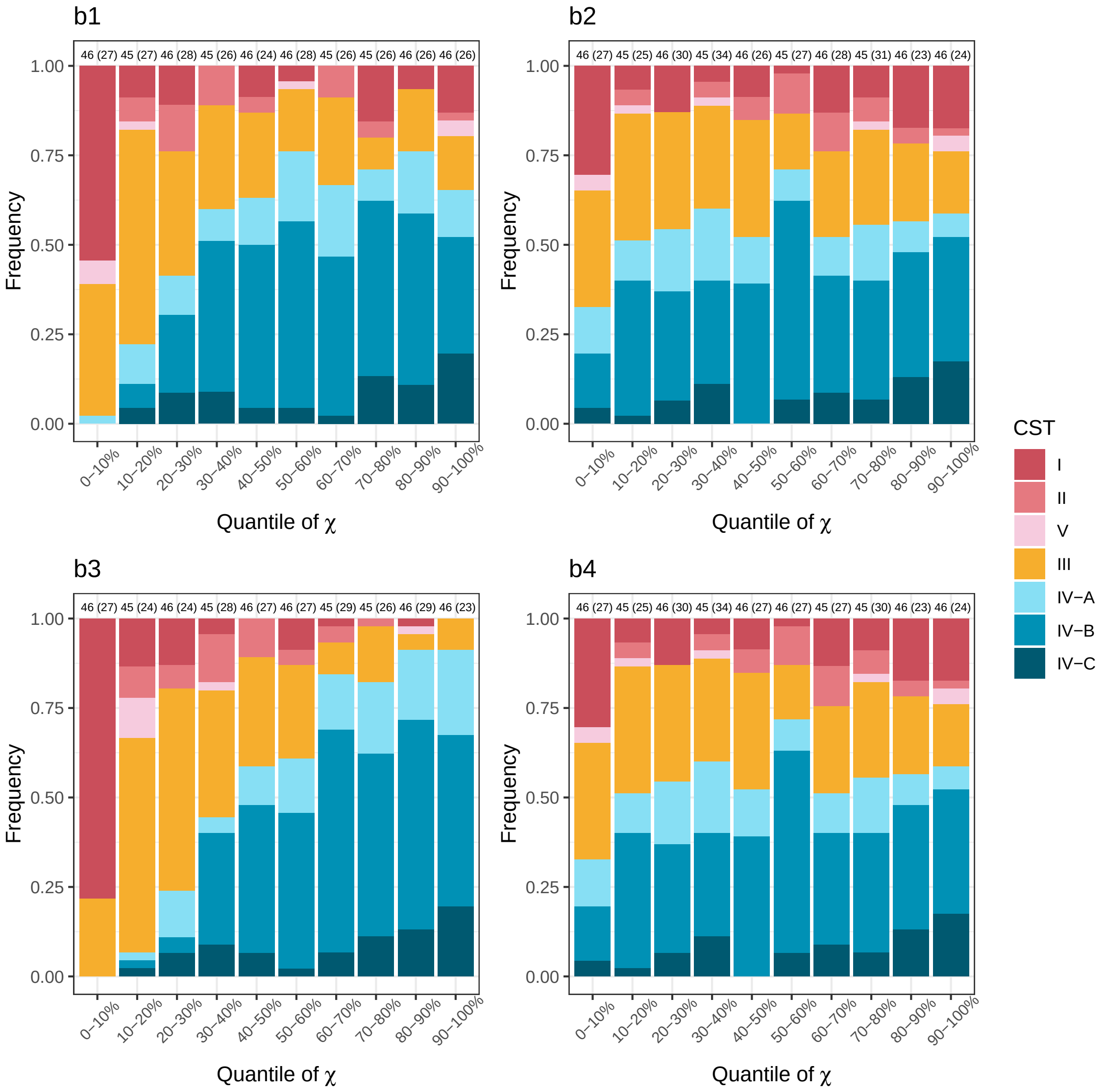


Figure S4: The frequencies of CST as a function of quantiles of empirically estimated $\chi$ (i.e., the proportion of supplied resource private to CST IV) assuming different half-saturation constants, $\kappa$ and $\theta$. The panel labels (i.e., b1, b2, b3, b4) correspond to the position in the parameter space indicated in Fig. S3.b with b1 indicating the default parameter values. The numbers on top of each bar indicate the numbers of samples (and women in parentheses), respectively.

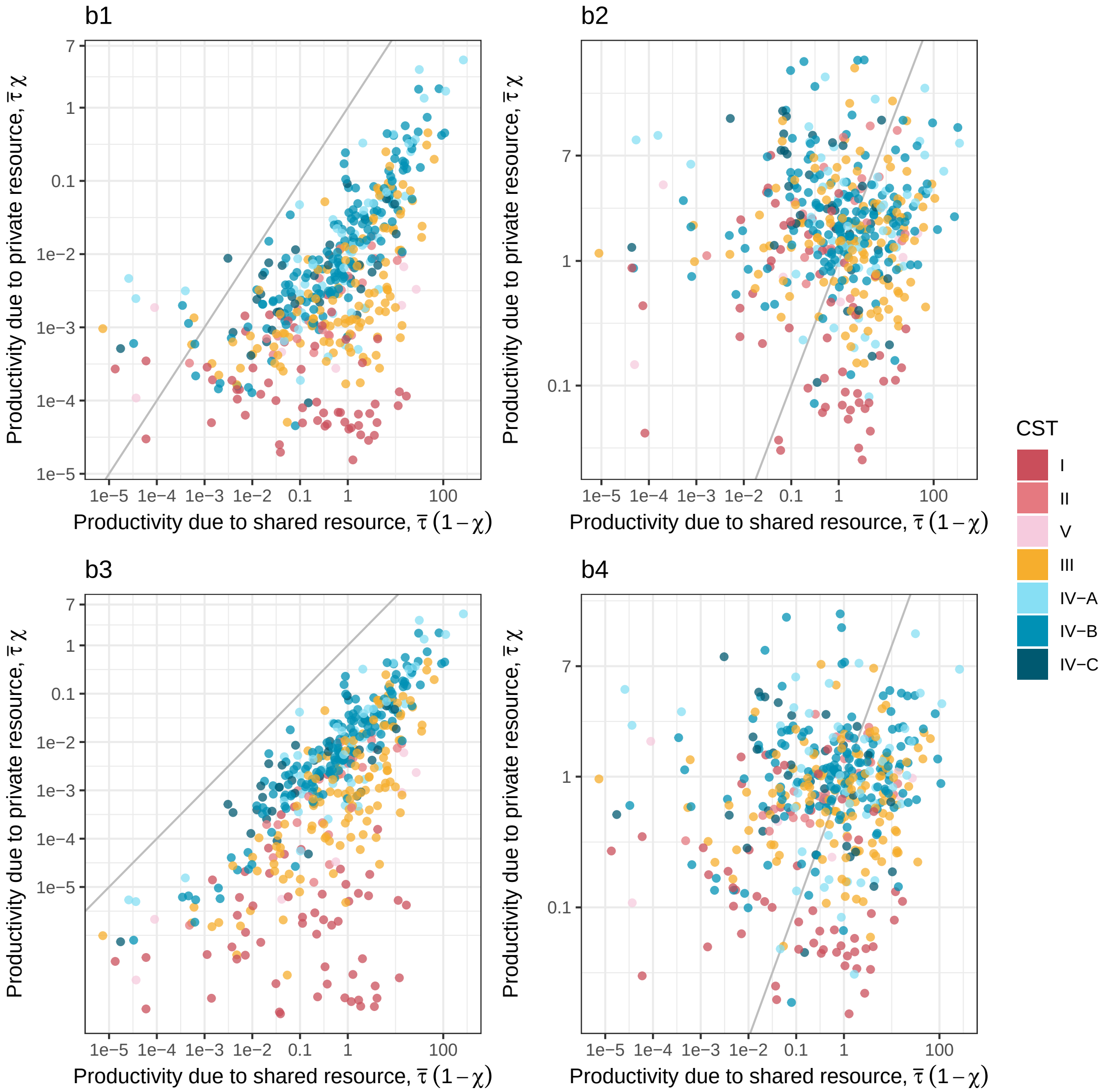


Figure S5: CST classification as a function of the productivity due to shared (x-axis) and private resources (y-axis) assuming different half-saturation constants, $\kappa$ and $\theta$. The panel labels (i.e., b1, b2, b3, b4) correspond to the position in the parameter space indicated in Fig. S3.b, with b1 indicating the default parameter values. The grey diagonal line indicates an equal supply of the two resource types.

### References


**Anukam KC, Reid G. 2008**. Effects of metronidazole on growth of *Gardnerella vaginalis* atcc 14018, probiotic *Lactobacillus rhamnosus* gr-1 and vaginal isolate *Lactobacillus plantarum* kca. *Microbial Ecology in Health and Disease* **20**:48–52.

**Argentini C, Fontana F, Alessandri G, Lugli GA, Mancabelli L, Ossiprandi MC, et al. 2022**. Evaluation of modulatory activities of *Lactobacillus crispatus* strains in the context of the vaginal microbiota. *Microbiology Spectrum* **10**:e02733–21.

**Atassi F, Brassart D, Grob P, Graf F, Servin AL. 2006**. *Lactobacillus* strains isolated from the vaginal microbiota of healthy women inhibit *Prevotella bivia* and *Gardnerella vaginalis* in coculture and cell culture. *FEMS Immunology & Medical Microbiology* **48**:424–432.

**Brown CT, Olm MR, Thomas BC, Banfield JF. 2016**. Measurement of bacterial replication rates in microbial communities. *Nature Biotechnology* **34**:1256–1263.

**Droop MR. 1968**. Vitamin b12 and marine ecology. iv. the kinetics of uptake, growth and inhibition in monochrysis lutheri. *Journal of the Marine Biological Association of the United Kingdom* **48**:689–733.

**France M, Alizadeh M, Brown S, Ma B, Ravel J. 2022**. Towards a deeper understanding of the vaginal microbiota. *Nature Microbiology* **7**:367–378.

**Gajer P, Brotman RM, Bai G, Sakamoto J, Schütte UM, Zhong X, et al. 2012**. Temporal dynamics of the human vaginal microbiota. *Science Translational Medicine* **4**:132ra52–132ra52.

**Harwich MD, Alves JM, Buck GA, Strauss JF, Patterson JL, Oki AT, et al. 2010**. Drawing the line between commensal and pathogenic *Gardnerella vaginalis* through genome analysis and virulence studies. *BMC Genomics* **11**:1–12.

**Langmead B, Salzberg SL. 2012**. Fast gapped-read alignment with bowtie 2. *Nature methods* **9**:357–359.

**Lee CY, Cheu RK, Lemke MM, Gustin AT, France MT, Hampel B, et al. 2020**. Quantitative modeling predicts mechanistic links between pre-treatment microbiome composition and metronidazole efficacy in bacterial vaginosis. *Nature Communications* **11**:1–12.

**O'Hanlon DE, Moench TR, Cone RA. 2013**. Vaginal ph and microbicidal lactic acid when lactobacilli dominate the microbiota. *PLOS One* **8**:e80074.

**Van Der Veer C, Hertzberger RY, Bruisten SM, Tytgat HL, Swanenburg J, de Kat Angelino-Bart A, et al. 2019**. Comparative genomics of human *Lactobacillus crispatus* isolates reveals genes for glycosylation and glycogen degradation: implications for in vivo dominance of the vaginal microbiota. *Microbiome* **7**:1–14.
